# Genomic resources for *Macadamia tetraphylla* and an examination of its historic use as a crop resource in Hawaii

**DOI:** 10.1101/2023.12.22.573098

**Authors:** Bjarne Bartlett, Alyssa Cho, Daniel Laspisa, Michael A. Gore, Michael Kantar

## Abstract

*Macadamia tetraphylla* is a wild relative of the economically valuable crop *Macadamia integrifolia*. Genomic knowledge of crop wild relatives is central to determining their possible role in breeding programs to mitigate biotic and abiotic stress in the future. The goal of this project was to develop a genomic resource for macadamia agriculture in Hawai‘i through constructing a transcriptome of *M. tetraphylla* and testing for hybridity in University of Hawai‘i at Mānoa breeding material. The transcriptome assembly of *M. tetraphylla* revealed large differences in gene expression attributable to tissue type. Advanced breeding lines (HI862 and HI879) appear to be hybridized with the crop wild relative *M. tetraphylla*. Additionally, a putative *M. tetraphylla* tree sampled from a remnant orchard planting at the Waimānalo research station on Oahu did not match anecdotal accounts of the orchard as it appeared to be of hybrid ancestry.

## Introduction

Macadamia nut (*Macadamia integrifolia*; 2*n* = 2*x* =28) is an important global crop originating in Australia (Mueller, 1858; Lin et al., 2022]. Recently, there has been increased interest in developing genomic resources for this crop (e.g. genome – Nock et al., 2020) due to its importance as a high-value, nutrient-dense food (Rengel et al., 2015;Navarro et al., 2016) that has undergone relatively little commercial breeding. In Hawai‘i alone, there has been an increase in resources dedicated to the crop due to its large land uses (17,100 acres in 2019) and high value ($42 million USD) [6]. These efforts have identified agronomic problems in the current orchards that can be attributed to the need for genetic improvement (Gutierrez-Coarite, et al., 2021). There is, however, limited genetic diversity within University of Hawai‘i *M. integrifolia* germplasm (Hardner, 2016). Many factors threaten the livelihoods of the Hawai‘i macadamia industry including pest and disease pressure; however, there is no adapted *M. integrifolia* tolerance for the most important pressures, requiring the use of a different donor species.

In species with limited genetic diversity, the field of plant breeding commonly uses related species as a donor to improve specific traits (Castaneda-Alvarez et al., 2016). In fact, the genetic diversity from crop wild relatives have been valued at over $100 billion U.S. dollars annually [(PwC, 2013]. The most important relative of *M. integrifolia* is *Macadamia tetraphylla* (2*n* = 2*x* = 28), a species that is occasionally cultivated but more frequently used as a donor for useful biotic stress tolerance traits (Mueller, 1858; Nin et al., 2022). To best operationalize the use of unadapted plant material for specific localities, genomic information is required (Wambugu and Henry, 2022). While there is a history of breeding in Hawai‘i gaps in staffing have caused records to be in doubt; therefore, there is also a need to reveal genetic ancestry in current promising breeding lines. Recently, a genome of *M. tetraphylla* was released (Nin et al., 2022), providing a useful resource to characterize Hawaiian *M. tetraphylla*. Therefore, the goals of this study were to i) characterize the *M. tetraphylla* transcriptome and ii) explore the ancestry of promising University of Hawai‘i at Mānoa breeding lines.

## Materials and Methods

A transcriptome assembly of *M. tetraphylla* was constructed from existing public data along with new RNAsequencing (RNAseq) data generated from wild-collected, feral *M. tetraphylla* and putative hybrid breeding lines in Hawai‘i with RNA from leaf tissue extracted using RNAeasy Plant Mini Kit (Qiagen, Valencia, California). RNA was then sent to Novogene Bioinformatics Technology Co., Ltd., for sequencing, which was conducted on an Illumina NovaSeq with 150 bp paired-end reads. Next-generation sequencing data (PRJNA587821) was obtained for *M. tetraphylla* for five different tissue types and assembled using Trinity (Haas et al., 2013). Coding regions within transcripts were identified with TransDecoder V.5.5.0 (Haas et al., 2012). Annotation of transcripts was done with BLASTX (Camacho et al., 2009) using the UniProt database to identify homologous sequences (UniProt Consortium, 2021). The HMMER (Eddy, 2011) and PFAM (Mistry et al., 2021) databases were used to identify protein domains. Data for all analyses are reported in **Table 1**. A read set was generated using combined RNAseq data from 5 tissues (bark, proteoid root, flowering inflorescence, young inflorescence, and leaves; SRR10424518-SRR10424522) that were trimmed, and quality checked using the same methods as for the Trinity assembly. The combined read set was mapped to the *M. tetraphylla* genome assembly (ASM2298504v1) using TopHat2 (default parameters). The resulting alignment file was filtered for unmapped reads (samtools view -F 0×04), sorted, and indexed using samtools. Expression of transcripts was determined by assessing FPKM (fragments per kilobase million), a transcript was considered expressed if FPKM > 10. Annotations were generated from the sorted alignment file using Cufflinks 2.2.1 (default parameters), which generated the GTF file used in the analysis. The alignment file and annotations were manually inspected in IGV for validation. The tool sppIDer (Langdon et al., 2018) was used to assess hybridity.

**Table 1:** Overview of data files.

| Label | Name of data file | File type | Data repository and identifier |
| --- | --- | --- | --- |
| Data set 1 | SRR10424522_leaves.fasta | .fasta | NCBI Sequence Read Archive:<br>https://identifiers.org/ncbi/insdc.sra:SRR10424522<br>Figshare:<br>https://doi.org/10.6084/m9.figshare.21084853.v1<br>(Bartlett et al., 2022a)<br>Zenodo:<br>https://doi.org/10.5281/zenodo.7135297 (Bartlett et al., 2022b) |
| Data set 2 | SRR10424521_younginflorescence.fasta | .fasta | NCBI Sequence Read Archive:<br>https://identifiers.org/ncbi/insdc.sra:SRR10424521<br>Figshare:<br>https://doi.org/10.6084/m9.figshare.21084853.v1<br>(Bartlett et al., 2022a)<br>Zenodo:<br>https://doi.org/10.5281/zenodo.7135297 [21] |
| Data set 3 | SRR10424520_floweringinflorescence.fasta | .fasta | NCBI Sequence Read Archive:<br>https://identifiers.org/ncbi/insdc.sra:SRR10424520<br>Figshare:<br>https://doi.org/10.6084/m9.figshare.21084853.v1<br>(Bartlett et al., 2022a)<br>Zenodo:<br>https://doi.org/10.5281/zenodo.7135297 [21] |
| Data set 4 | SRR10424519_proteoidroot.fasta | .fasta | NCBI Sequence Read Archive:<br>https://identifiers.org/ncbi/insdc.sra:SRR10424519<br>Figshare:<br>https://doi.org/10.6084/m9.figshare.21084853.v1<br>(Bartlett et al., 2022a)<br>Zenodo:<br>https://doi.org/10.5281/zenodo.7135297 [21] |
| Data set 5 | SRR10424518_bark.fasta | .fasta | NCBI Sequence Read Archive:<br>https://identifiers.org/ncbi/insdc.sra:SRR10424518<br>Figshare:<br>https://doi.org/10.6084/m9.figshare.21084853.v1<br>(Bartlett et al., 2022a)<br>Zenodo:<br>https://doi.org/10.5281/zenodo.7135297 [21] |
| Data set 6 | MT1_879.vcf.gz | .vcf.gz | Figshare:<br>https://doi.org/10.6084/m9.figshare.21084853.v1<br>(Bartlett et al., 2022a) |
|  |  |  | Zenodo:<br>https://doi.org/10.5281/zenodo.7135297 (Bartlett et al., 2022b)<br>European Variation Archive:<br>https://identifiers.org/ebi/bioproject:PRJEB56438) |
| Data set 7 | MT5_OT.vcf.gz | .vcf.gz | Figshare:<br>https://doi.org/10.6084/m9.figshare.21084853.v1 (Bartlett et al., 2022a)<br>Zenodo:<br>https://doi.org/10.5281/zenodo.7135297 (Bartlett et al., 2022b)<br>European Variation Archive:<br>https://identifiers.org/ebi/bioproject:PRJEB56438 ) |
| Data set 8 | MT9_OTPH.vcf.gz | .vcf.gz | Figshare:<br>https://doi.org/10.6084/m9.figshare.21084853.v1 (Bartlett et al., 2022a)<br>Zenodo:<br>https://doi.org/10.5281/zenodo.7135297 (Bartlett et al., 2022b)<br>European Variation Archive:<br>https://identifiers.org/ebi/bioproject:PRJEB56438 |
| Data set 9 | MT10_862.vcf.gz | .vcf.gz | Figshare:<br>https://doi.org/10.6084/m9.figshare.21084853.v1 (Bartlett et al., 2022a)<br>Zenodo:<br>https://doi.org/10.5281/zenodo.7135297 (Bartlett et al., 2022b)<br>European Variation Archive:<br>https://identifiers.org/ebi/bioproject:PRJEB56438 |
| Data set 10 | MT1_1.fastq.gz<br>MT1_2.fastq.gz | .fq.gz | NCBI Sequence Read Archive:<br>https://identifiers.org/ncbi/insdc.sra:SRR21236275 |
| Data set 11 | MT10_1.fq.gz<br>MT10_2.fq.gz | .fq.gz | NCBI Sequence Read Archive:<br>https://identifiers.org/ncbi/insdc.sra:SRR21236272 |
| Data set 12 | MT9_1.fq.gz<br>MT9_2.fq.gz | .fq.gz | NCBI Sequence Read Archive:<br>https://identifiers.org/ncbi/insdc.sra:SRR21236273 |
| Data set 13 | MT5_1.fq.gz<br>MT5_2.fq.gz | .fq.gz | NCBI Sequence Read Archive:<br>https://identifiers.org/ncbi/insdc.sra:SRR21236274 |

## Results

### Differential gene expression across tissue type

The *de novo M. tetraphylla* transcriptomes showed a high level of completeness (**Figure 1a**) using Benchmarking Universal Single-Copy Orthologous (BUSCO) for bark (97.65% complete), proteoid root (98.43% complete), flowering inflorescence (98.43% complete), young inflorescence (99.61% complete), and leaves (98.82% complete). For validation and annotation, the assembled sequence was compared to the KEGG and PFAM databases (Mistry et al., 2021; Ogata et al., 1998). Similar to a previous report (Niu et al., 2022), a total of 46,967 transcripts were called representing 34,444 unigenes and 12,523 isoforms. As expected, there was tissue specific expression of transcripts and pathways (FPKM > 10 **Figure 1b, c**).

**Figure 1.**
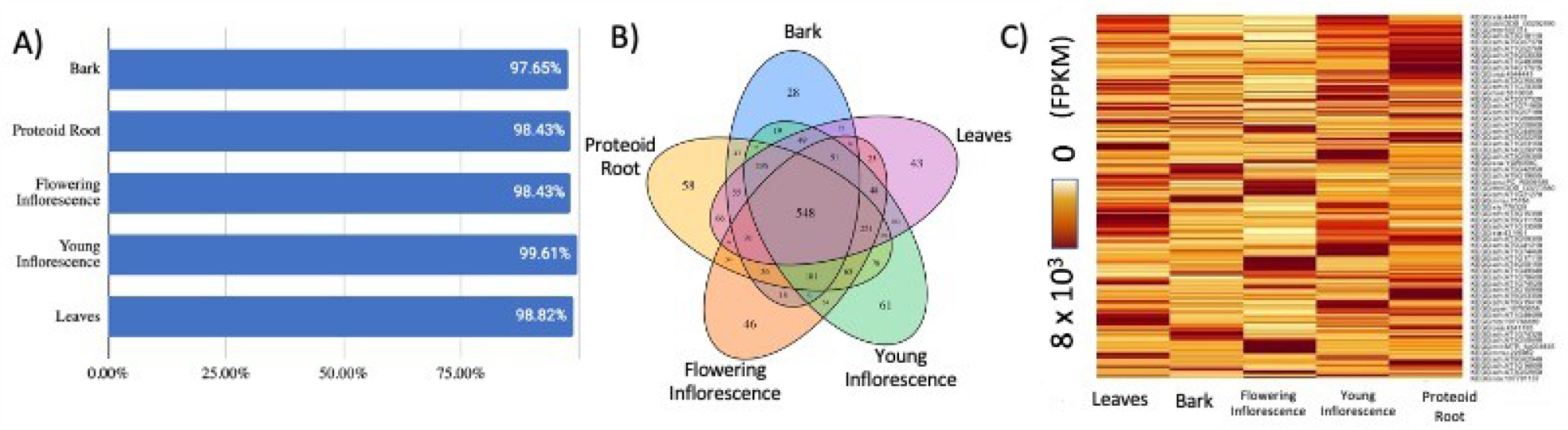
A) The de novo M. tetraphylla transcriptomes showed high completeness based on Benchmarking Universal Single-Copy Orthologous (BUSCO) quality assessments. B) Number of genes expressed in different tissues by KEGG Pathway in M. tetraphylla. Total number of genes with FPKM > 10 across proteoid root, flowering inflorescence, young inflorescence, and leaves. C) Gene expression of KEGG pathways represented as FPKM (fragments per kilobase million) in M. tetraphylla for leaves, bark, flowering inflorescence, young inflorescence, and proteoid root.

### Hybridity of Hawai‘i Collected Individuals

Combining the information from the *de novo* assembly and the previous sequence showed that the new assembly has potential to inform *M. integrifolia* breeding. We identified control samples from previously published data to calibrate putative hybridization (**Figure 2A, B**). We sampled a tree (Oahu Putative Hybrid from the Tantalus Lookout Puu Ualakaa State Park - **Figure 2C**) from a remnant orchard planting that showed ancestry percentages supporting a hybrid origin. However, its percentage of *M. integrifolia* ancestry was estimated to be ∼35%, suggesting that this putative hybrid is an F_1_ with some segregation distortion or a late generation hybrid. Surprisingly, the *M. tetraphylla* from a remnant planting at Waimānalo (**Figure 2E**) also was inferred to be a hybrid, again likely an F_1_ or a later generation hybrid. Additionally, we found close to the expected ancestry percentage (25%) for two putative advanced BC_1_ ((*M. tetraphylla* x *M. integrifolia*) x *M. integrifolia*)) breeding lines (HI862 - 19.6% and HI879 - 24.8%) from the UH Mānoa breeding program (**Figure 2D and F; Supplemental Table 1**). However, the 13.4% *M. tetraphylla* in the *M. integrifolia* control (Figure 2B) may instead indicate that the breeding lines are BC_2_ rather than BC_1_. Introgressions from *M. tetraphylla* appeared to be consistent within HI862 and HI879, occurring on chromosome 2, 5, and 9 (**Supplemental Figure 1**).

**Figure 2.**
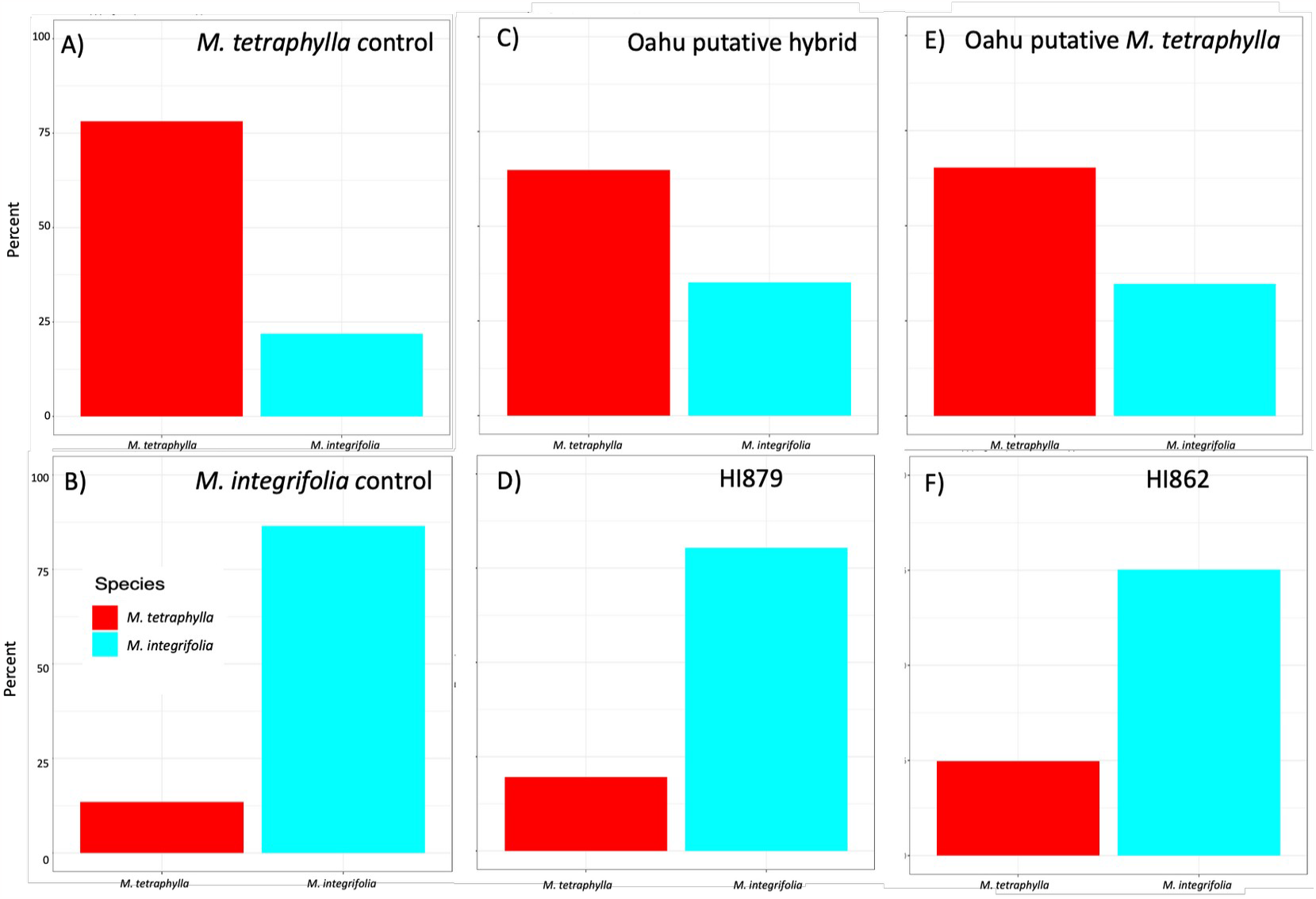
Hybridity of Hawai‘i collected individuals. A) M. tetraphylla control (SRR10424522), B) M. integrifolia control (PRJNA593881), C) Putative hybrid collected from a remnant orchard planting in Tantalus Lookout Puu Ualakaa State Park on Oahu, D) Putative hybrid breeding line from UH Mānoa, E) M. tetraphylla collected from the UH Mānoa Waimānalo research station on Oahu, and F) Putative hybrid breeding line from UH Mānoa.

## Discussion

It is now a routine procedure to construct genomic resources for non-model crop species and their relatives (Wambugu and Henry, 2022). This is an important step in increasing the efficiency of interspecific breeding, especially in species that have long generation time (Valle-Echevarria et al., 2021). Transcriptomes provide insight into gene function and are a potentially rich source of information on crop relatives. Here, we showed considerable tissue-specific expression across functional pathways. Furthermore, we found evidence of hybridization not only in breeding lines, but also unexpectedly for a putative *M. tetraphylla* in remnant orchard material at the Waimānalo research station on Oahu. Although the breeding lines had the expected levels of *M. tetraphylla* when compared to pedigrees, additional generations of backcrossing to *M. integrifolia* could be needed for production in commercial Hawaiian orchards. There are some limitations associated with our data, (1) short read data (∼150bp) has limited ability to explore multiple isoforms, (2) we only have single control for each sample, and (3) the limited sampling in remnant seed orchards prevented a more complete characterization of potential genetic variation for introgressions. Wild relatives of crops have huge potential value, but in order to maximize the ability to use them they will need to be extensively characterized.

## Abbreviations

CWR: crop wild relative

## Declarations

### Availability of data and materials

The data described here can be freely and openly accessed at: https://doi.org/10.6084/m9.figshare.21084853.v1 -- (Transcriptome Leaf of M. tetraphylla, Transcriptome young inflorescence of M. tetraphylla, Transcriptome flowering inflorescence of M. tetraphylla, Transcriptome proteoid root of M. tetraphylla, Transcriptome bark of M. tetraphylla, VCF file of M. integrifolia × M. tetraphylla hybrid Hawaii 879, VCF file of Oahu M. tetraphylla, VCF file of Oahu putative M. integrifolia × M. tetraphylla hybrid, VCF file of M. integrifolia × M. tetraphylla hybrid Hawaii 862 – [20]) and at Zenodo. https://doi.org/10.5281/zenodo.7135297 -- (Transcriptome Leaf of M. tetraphylla, Transcriptome young inflorescence of M. tetraphylla, Transcriptome flowering inflorescence of M. tetraphylla, Transcriptome proteoid root of M. tetraphylla, Transcriptome bark of M. tetraphylla, VCF file of M. integrifolia × M. tetraphylla hybrid Hawaii 879, VCF file of Oahu M. tetraphylla, VCF file of Oahu putative M. integrifolia × M. tetraphylla hybrid, VCF file of M. integrifolia × M. tetraphylla hybrid Hawaii 862 [21]).DNA from MT1, MT5, MT9 and MT10 upon reasonable request.

The variant data for this study have been deposited in the European Variation Archive (EVA) at EMBL-EBI under accession number PRJEB56438 (https://identifiers.org/ebi/bioproject:PRJEB56438 – [22]

*Raw sequence data are available at:*

Yunnan Institute of Tropical Crops: RNA seq of Macadamia tetraphylla: leaves 2019; NCBI Sequence Read Archive: https://identifiers.org/ncbi/insdc.sra:SRR10424522 [23]

Yunnan Institute of Tropical Crops: RNA seq of Macadamia tetraphylla9: young inflorescence; NCBI Sequence Read Archive: https://identifiers.org/ncbi/insdc.sra:SRR10424521 [24]

Yunnan Institute of Tropical Crops: RNA seq of Macadamia tetraphylla: flowering inflorescence 2019: NCBI Sequence Read Archive: https://identifiers.org/ncbi/insdc.sra:SRR10424520 [25]

Yunnan Institute of Tropical Crops: RNA seq of Macadamia tetraphylla: proteoid root 2019; NCBI Sequence Read Archive: https://identifiers.org/ncbi/insdc.sra:SRR10424519 [26]

Yunnan Institute of Tropical Crops: RNA seq of Macadamia tetraphylla: bark 2019; NCBI Sequence Read Archive: https://identifiers.org/ncbi/insdc.sra:SRR10424518 [27]

University of Hawaii: Hawaii 879 2022; NCBI Sequence Read Archive: https://identifiers.org/ncbi/insdc.sra:SRR21236275 [28]

University of Hawaii: Hawaii 862 2022; NCBI Sequence Read Archive: https://identifiers.org/ncbi/insdc.sra:SRR21236272 [29]

University of Hawaii: Oahu putative hybrid Tantalus 2022; NCBI Sequence Read Archive: https://identifiers.org/ncbi/insdc.sra:SRR21236273 [30]

University of Hawaii: Oahu Tetraphylla Waimanalo 2022; NCBI Sequence Read Archive: https://identifiers.org/ncbi/insdc.sra:SRR21236274 [31]

## Authors’ contributions

BB: Analysis, data generation, drafting, revision, AC: Conceptualization, writing/revision, DL: Analysis, writing/revision, MK: Conceptualization, writing/revision, MG: Conceptualization, writing/revision

## Acknowledgements

We would like to thank Cornell University for supporting the sabbatical of Dr. Michael A. Gore so he could contribute to this manuscript.

