## Supplemental Figure 1 for "Genomic resources for *Macadamia tetraphylla* and an examination of its historic use as a crop resource in Hawaii"

**Supplemental Figure 1.** Average depth of coverage across the entire genome for HI879 and HI862. Red indicates *M. tetraphylla* depth and teal indicates *M. integrifolia* depth. Vertical dotted lines represent chromosomes. Horizontal lines show bins demarcated by the local minima of the coverage distribution, those places at the extreme ends of the empirical distribution are considered locations of introgression (read depth > 40).

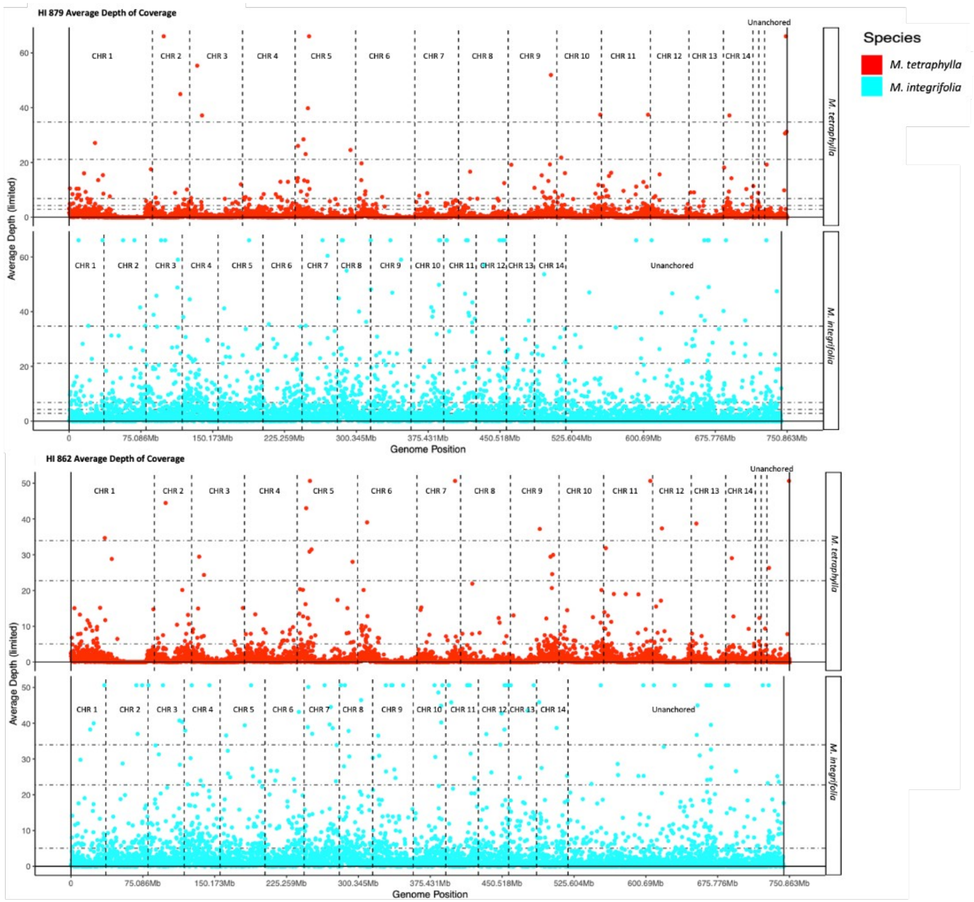
