## Supplemental Table 1 for "Genomic resources for *Macadamia tetraphylla* and an examination of its historic use as a crop resource in Hawaii"

| | MT-1 -HI 879 | | | MT-10-Hawaii 862 | | | MT-5-Oahu Tetraphylla | | | MT-9-Oahu putative hybrid | | | Control MacInt | | | Control MacTet | | |
|---|---|---|---|---|---|---|---|---|---|---|---|---|---|---|---|---|---|---|
| | Unmapped | MacTet | MacInt | Unmapped | MacTet | MacInt | Unmapped | MacTet | MacInt | Unmapped | MacTet | MacInt | Unmapped | MacTet | MacInt | Unmapped | MacTet | MacInt |
| Number Reads | | 54262791 | | | 54241998 | | | 46657000 | | | 43205846 | | | 460617384 | | | 51773462 | |
| Number Mapped Reads | | 53873335 | | | 54158604 | | | 45630572 | | | 42479302 | | | 459150448 | | | 42861472 | |
| Percent Unmapped Reads | | 48.67% | | | 50.39% | | | 48.59% | | | 49.25% | | | 16.83% | | | 58.02% | |
| Average MQ | | 18.80650706 | | | 17.59945944 | | | 18.44314973 | | | 17.78028851 | | | NA | | | 14.92664823 | |
| Median MQ | | 6 | | | 0 | | | 6 | | | 3 | | | 49 | | | 0 | |
| Total mapped reads | 389456 | 17512904 | 36360431 | 83394 | 19408531 | 34750073 | 1026428 | 26324256 | 19306316 | 726544 | 24602802 | 17876500 | 1466936 | 78855673 | 380294775 | 8911990 | 27822186 | 15039286 |
| % of all reads | 0.72% | 32.27% | 67.01% | 0.15% | 35.78% | 64.06% | 2.20% | 56.42% | 41.38% | 0.0168 | 0.5694 | 0.4138 | 0.32% | 17.12% | 82.56% | 17.21% | 53.74% | 29.05% |
| % Nonzero mapped reads | 0% | 19.60% | 80.40% | 0% | 24.82% | 75.18% | 0% | 64.82% | 35.18% | 0 | 0.6528 | 0.3472 | 0% | 13.49% | 86.51% | 0.00% | 78.12% | 21.88% |
| All average MQ | 0 | 9.48 | 23.5 | 0 | 10.31 | 21.71 | 0 | 20.71 | 16.33 | 0 | 19.84 | 15.67 | 0 | 26.07 | 0 | 0 | 21.22 | 12.12 |
| NonZero average MQ | 0 | 30.41 | 38.16 | 0 | 29.95 | 37.3 | 0 | 35.07 | 37.37 | 0 | 34.11 | 36.79 | 0 | 39.79 | 0 | 0 | 34.78 | 38.34 |
| All Median MQ | 0 | 0 | 11 | 0 | 0 | 10 | 0 | 10 | 0 | 0 | 9 | 0 | 0 | 27 | 57 | 0 | 10 | 0 |
| Nonzero Median MQ | 0 | 27 | 40 | 0 | 27 | 40 | 0 | 33 | 40 | 0 | 27 | 40 | 0 | 40 | 60 | 0 | 34 | 40 |

**Supplemental Table 1: Average depth of coverage across the entire genome for HI879 and HI862. Red indicates M. tetraphylla depth and teal indicates M. integrifolia depth. Vertical dotted lines represent chromosomes. Horizontal lines show bins demarcated by the local minima of the coverage distribution; those places at the extreme ends of the empirical distribution are considered locations of introgression (read depth > 40).**
